# Male-specific CREB signaling in the hippocampus controls spatial memory deficits in a mouse model of autism and intellectual disability

**DOI:** 10.1101/249649

**Authors:** Marta Zamarbide, Adele Mossa, Molly K. Wilkinson, Heather L. Pond, Adam W. Oaks, M. Chiara Manzini

## Abstract

**Background:** The prevalence of neurodevelopmental disorders is biased towards males with male: female ratios of 2:1 in intellectual disability (ID) and 4:1 in autism spectrum disorder (ASD). However, the molecular mechanisms of such bias remain unknown. While characterizing a mouse model for loss of the signaling scaffold *coiled-coil and C2 domain containing 1A* (CC2D1A), which is mutated in ID and ASD, we identified biochemical and behavioral differences between males and females, and explored whether CC2D1A controls male-specific intracellular signaling.

**Methods:** CC2D1A is known to regulate phosphodiesterase 4D (PDE4D). We tested for activation PDE4D and downstream signaling molecules such as CREB in the hippocampus of *Cc2d1a*-deficient mice. We then performed behavioral studies in females to analyze learning and memory, social interactions, anxiety and hyperactivity. Finally, we targeted PDE4D activation with a PDE4D inhibitor to define how changes in PDE4D and CREB activity affect behavior in males and females.

**Results:** We found that in *Cc2d1a*-deficient males PDE4D is hyperactive leading to a reduction in CREB signaling, but this molecular deficit is not present in females. *Cc2d1a*-deficient females only show impairment in novel object recognition, and no other cognitive and social deficits that have been found in males. Restoring PDE4D activity using an inhibitor rescues male-specific cognitive deficits, but has no effect on females.

**Conclusions:** Our findings show that CC2D1A regulates intracellular signaling in a male-specific manner in the hippocampus leading to male-specific behavioral deficits. We propose that male-specific signaling mechanisms are involved in establishing sex bias in neurodevelopmental disorders.

## INTRODUCTION

Developmental disabilities are more prevalent in males with male: female ratios ranging between 1.5-2:1 in intellectual disability (ID) and 4-10:1 in autism spectrum disorders (ASD) (1–3). However, the molecular mechanisms underlying sex bias in neurodevelopmental disorders remain unknown. The male and female brain show differences in normal subjects beyond behaviors that are commonly considered sex-specific, such as mating and aggression (4). Non-invasive brain imaging studies in humans have shown that male and female brains have different patterns of connection (5), and comparison of activation of brain areas involved in social cognition in children with ASD found deficits only in affected boys, and not in girls (6). In parallel, studies testing learning and memory in rodents showed that lesions in certain brain regions result in different deficits in males and females (7,8). For example, in some spatial learning task males may rely on hippocampal circuits, while females predominantly use the striatum (9). Similarly, in context fear memory retrieval more hippocampal neurons are activated in males, while the basal amygdala is more active in females (10). These findings suggest that males and females may use different cellular and molecular strategies to encode information for the same behaviors.

In particular, intracellular signaling differs in males and females and is a candidate for differentially controlling behavioral outputs. Activity of cAMP response element binding protein (CREB) downstream of protein kinase A (PKA) and mitogen-activated protein (MAPK/ERK) kinases is differentially regulated by estrogen in the hippocampus of males and females throughout development (11,12). CREB activity in the hippocampus is critical for memory formation and retrieval (13,14), and studies in primary hippocampal neurons showed that CREB is primarily controlled by estrogen receptor activation in female neurons, and not in males (11). A study in a mouse model of 16p11.2 deletion syndrome found male specific hyperactivity in MAPK3/ERK1 kinase in the striatum correlating with male-specific striatal learning deficits (15), suggesting a possible link between sex-specific signaling and behavior.

Here, we introduce the signaling scaffold *coiled-coil and C2 domain containing 1A* (CC2D1A), as a regulator of sex-specific intracellular signaling in the male hippocampus controlling spatial memory acquisition. Biallelic mutations in *CC2D1A* cause a spectrum of neurodevelopmental conditions in humans with fully penetrant ID and variably penetrant ASD, ADHD, seizures, and aggressive behavior (16–19). While global *Cc2d1a* knock-out pups die at birth due to respiratory deficits, we found that conditional removal of this gene in the cortex and hippocampus leads to cognitive and social deficits, hyperactivity, anxiety, and self-injury in males (20). *In vitro* studies in mouse embryonic fibroblasts and hippocampal neurons from *Cc2d1a* global KO mice have shown that CC2D1A binds to phosphodiesterase (PDE4D), an enzyme involved in cAMP degradation, and regulates PDE4D activity and subcellular distribution (21,22). When CC2D1A is lost, PDE4D is phosphorylated and becomes hyperactive, leading to decreased intracellular levels of cAMP and a reduction in phosphorylation of CREB via PKA (22). We wondered whether PDE4D and CREB function was perturbed in the *Cc2d1a*-deficient hippocampus and whether these changes caused the spatial cognitive deficits observed in the mice. During our studies, we found that only *Cc2d1a*-deficient males show disruptions in PDE4D signaling, which correlate with male-specific spatial memory deficits. We that used a PDE4D inhibitor to rescue these deficits confirming that restoring signaling directly affects behavior. Females do not show these molecular and behavioral impairments. Our results show that CC2D1A regulates CREB signaling in a sex-specific manner in males, revealing a novel mechanism controlling male-specific intracellular signaling and behavior.

## METHODS AND MATERIALS

### Animals

All animal care and use was in accordance with institutional guidance and approved by the Institutional Animal Care and Use Committee of The George Washington University. The *Cc2d1a* conditional knock-out (cKO) mouse line was generated by crossing Cc2d1a-flx mice (20) with a CamKIIa-cre mouse line driving Cre recombinase expression under the *CamKIIa* promoter (Stock 005359, Jackson Laboratories) (23). All animals are fully backcrossed on a C57BL/6 background for at least 6 generations. Since the CamKIIa-cre transgene can be occasionally activated in male germ cells leading to germline transmission (24), crosses were conducted between homozygous Cc2d1a-flx males and double heterozygous Cc2d1a-flx/CamKIIa-cre females. cKO males and female mice are viable and fertile as previously described (20). For genotyping, polymerase chain reaction (PCR) amplifications were performed on 1µL of proteinase K (New England Biolabs, Ipswich MA) digested tail DNA samples. PCR reactions (50µL) consisted of GoTaq Flexi buffer (Promega, Madison WI), 100µM dNTPs, 50µM each of forward and reverse primers (sequence available upon request), 1mM MgCl_2_, and 1.25 U GoTaq Flexi DNA polymerase (Promega, Madison WI), and were run with optimized reaction profiles determined for each genotype. A 25µL aliquot from each reaction was analyzed by gel electrophoresis on a 1.0% agarose gel for the presence of the desired band.

### Signaling analysis via Western blot and ELISA

Protein lysates were prepared from fresh, frozen mouse hippocampal tissue that was homogenized in a buffer containing Tris-HCl (50 mM), NaCl (100 mM), ethylenediaminetetraacetic acid (5 mM), magnesium chloride (2 mM), protease and phosphatase inhibitor mixture (Sigma-Aldrich, St. Louis MO) and extracted for 30 min with Triton X-100 (1%) and sodium dodecyl sulfate (0.1%). Lysates were cleared by centrifugation at 15,000x*g* for 20 min at 4°C then combined with one volume of Laemmli sample buffer (Bio-Rad, Hercules CA) containing 5% beta-mercaptoethanol and denatured by heating at 95°C for five min. Protein was separated by SDS-PAGE on 4-12% Bis-Tris gels (Thermo Scientific, Waltham MA) and transferred to Immobilon-FL (Millipore, Billerica MA) polyvinylidine fluoride membranes. Immunoblots were probed with primary antibodies at optimized concentrations: from FabGennix International (Frisco, TX) total PDE4D (PDE4D5-451AP, 1:500), pPDE4D Ser-190 (PPD4-440AP, 1:100); from Cell Signaling Technologies (Danvers, MA) total PKA (5842, 1:1,000), pPKA Thr-197 (5661, 1:500), total CREB (9197, 1:500), pCREB Ser-133 (9198, 1:250); from AbCam (Cambridge, MA) beta-actin (ab6276, 1:10,000), Cc2d1a (ab68302, 1:1,000). Primary antibodies were then labeled with infra-red fluorophore-conjugated secondary antibodies (LI-COR Biosciences, Lincoln NE) and imaged on an Odyssey Imager (LI-COR Biosciences, Lincoln NE).

Cyclic AMP (cAMP) levels were measured using the Cyclic AMP Complete ELISA Kit (AbCam, ab133051, Cambridge, MA) following instructions from the manufacturer. Briefly, flash frozen hippocampal tissue was extracted in 0.1M HCl and after HCl neutralization, mixed with anti-cAMP antibody and alkaline phosphatase for colorimetric detection in a Varioskan LUX multimode microplate reader (ThermoFisher Scientific, Waltham, MA).

### Behavioral tests

A standardized battery of behavioral testing was used for cKO animals at 3-4 months of age. Behavioral tests were performed in the Manzini lab behavior analysis suite in the George Washington University Animal Research Facility following a 60 min period of acclimatization. Initial characterization to test for of basic motor and somatosensory function was performed as described by Rogers (25): righting reflex, wire hang, gait analysis, tail pinch and visual reach. Cognitive and social function and other behaviors were tested in the open field test, novel object recognition test (26), Morris water maze (27) in one cohort, and 3-chamber social interaction test (28) in a second cohort of male and female littermate. Most results for the males cKO mice were published in Oaks *et al* (20). Behavioral analysis was performed via automated animal tracking using ANY-maze (Stoelting, Wood Dale IL).

### Motor function

Coordination, motor strength and vestibular function were tested by testing for the righting reflex and placing each mouse on its back. Ability to return to an upright position was timed. Motor strength was also tested via the wire hang test, by timing the latency to fall into an empty cage while hanging from a wire cage-top not higher than 18 cm. Finally, motor coordination was tested via gait analysis. Each mouse was placed into enough red non-toxic tempera paint to wet its paws, then was placed in front of narrow tunnel over white paper. Paw marks on the paper were analyzed for stride length and abnormalities in paw placement.

### Somatosensory function

Pain sensation was tested by pinching the tip of the tail with fine, ethanol-cleaned forceps. Reactions were categorized as either response or no response. Vision was tested by presenting a wire cage top while each mouse was being held by the base of the tail at a height of 18 cm over an open cage. Latency to reach for the wire cage top was measured.

### Novel Object Recognition Test

The Novel Object Recognition Test (NORT) (20,26) was performed in an 50x50X50cm plastic box (Stoelting, Wood Dale IL). The test consisted of three phases: habituation, training, and test. During the habituation phase the animals were exposed to the unfamiliar box for 30 min, then returned to the home cage while the box was being cleaned. For the training phase, the animal was placed back into box with two identical objects located in opposite corners 5 cm from the walls and allowed to explore for 10min. The animal was then returned to the home cage for 15 min and tested for short-term memory by placing a familiar object, identical to those used in the habituation phase, and an unfamiliar object at opposite corners of the test box. Exploration activity was again monitored for 10 min, with exploration defined as time spent actively observing or touching the object from within a radius of five cm. Cumulative time spent with each object was measured by video analysis using ANY-maze to determine the location of the animal’s nose relative to the objects in the enclosure. Preference for the novel object was defined as the ratio of the time spent with the novel object to the time spent with the familiar object. Mice were considered non-performers and removed from the analysis when they spent less than 2s interacting with the objects.

### Morris Water Maze

The Morris Water Maze (MWM) (20,27) apparatus was a 120x120cm round metal tub (Stoelting, Wood Dale IL), where distinct visual cues were placed at the cardinal points. The surface of the water was made opaque by adding white non-toxic paint and the temperature was maintained at 24 °C. Each daily test consisted of four independent trials, one at each cardinal point around the tub, with the mouse being placed in the water facing the wall of the tub. Each trial lasted until the mouse found the platform or up to 60s, whichever occurred first. Each animal completed two daily tests with a visible platform to learn that a platform is available to escape from the water, then performed five tests with a platform hidden under the water surface to memorize the platform location based on the cardinal cues on the walls of the apparatus. After the hidden platform trials a 60s probe trial was performed by removing the platform and measuring the time spent swimming over the correct platform location. Finally, two reversal tests were completed after changing the location of the platform to test for cognitive flexibility and the ability to learn a new platform location. Mice were considered non-performers and removed from the analysis when they refused to swim and floated in the water for 60 seconds. Only animals that completed all trials were included.

Learning strategy analysis was performed by analyzing all four cardinal point trials for each of the first three days of the hidden platform phase. Total swim path was tracked using AnyMaze and scored by an investigator blind to genotype as 1) thigmotaxis = swimming primarily by the walls of the tub; 2) random path = swimming throughout the tub with turns in all directions; 3) scanning = swimming the length of the tub in a zig-zag starting from the drop location; 4) circling = swimming in circles at a distance from the walls corresponding to the platform location; 5) directed path = swimming toward the general location of the platform with a few turns; 6) straight path = swimming directly to the platform (**Suppl. Fig.2A**). 1, 2 and 3 were considered motor strategies as they do not require a knowledge of the platform location, 4, 5 and 6 were considered place strategies as they imply that the animal remembers the distance and general position of the platform in the tub. Strategy usage was averaged across all animals.

### Open Field

The open field (OF) was conducted in an unfamiliar 50x50x50 cm plastic box (Stoelting, Wood Dale IL) and ambulatory activity was monitored by automated video tracking for 15 min. The arena was divided into two areas, an outer zone and a center zone (25x25 cm; 25% of total area). Total distance traveled, time spent in the periphery and the center, and entries into the center area were measured.

### Three-chamber social interaction test

The three-chambered test (28,29) was performed in a clear rectangular acrylic box (60x40 cm) divided into three chambers (40x20 cm) with small openings (10x5 cm) in the adjoining walls (Everything Plastic, Philadelphia PA). The test consisted of two phases, the habituation and the sociability phase. During the habituation phase, empty inverted wire cups (10 cm diameter) were placed in the center of the chambers at the ends. Each mouse was placed in the center chamber of the apparatus and allowed to explore the different chambers for five min. During the second phase, an unfamiliar mouse of the same sex as the tested mouse was placed under the wire cup in one of the side chambers. Experimental mice were allowed to explore for 10 min during the sociability phase. Total time spent in the Object (containing empty cup) and Mouse (with unfamiliar mouse under the cup) chambers was used to determine the social preference of each mouse tested, while the time sniffing within a 2-cm radius of the mouse-containing cup were recorded as measures of social approach and social interaction.

### Drug treatments

Animals were treated with 30µg/kg of GEBR-7b (Millipore, Burlington, MA) via intraperitoneal injection for 14 days prior to behavioral testing. Control animals were injected with saline only. Drug injections were continued every night during behavioral testing until the animals were sacrificed for tissue collection for Western blot analysis and ELISA, as described above.

## RESULTS

### CC2D1A controls signaling downstream of PDE4D in a male-specific manner

CC2D1A has been shown to bind directly to PDE4D and PKA (Fig.1A), and to regulate both PDE4D activity and subcellular localization in mouse embryonic fibroblasts and primary hippocampal neurons (21,22). It has been proposed that CC2D1A acts as a repressor of PDE4D, and prevents PDE4D phosphorylation via PKA, to control levels of cAMP within the cell (22). When cultured neurons lacking *Cc2d1a* are treated with forskolin to increase cAMP levels and activate CREB, hyperactive PDE4D reduces the effect of this drug leading to reduced PKA and CREB activation (21). Since we have recently found that conditional removal of *Cc2d1a* in the forebrain causes spatial memory deficits (20), which are often linked to reduced CREB activation in the hippocampus (13), we asked whether these deficits would be maintained in the adult hippocampus of *Cc2d1a*-deficient mice. We collected hippocampal tissue from 5 month old male and female control (homozygous Cc2d1a-LoxP, listed as WT) mice and Cc2d1a^CamKIIa-cre^(or cKO) mice, where *Cc2d1a* was removed postnatally in the hippocampus and cortex via Cre-recombinase expressed under the *CamKIIa* promoter as previously shown (20,23). We tested for PDE4D, PKA, and CREB phosphorylation via Western blot by determining the ratio of phosphorylated to total protein in hippocampal protein lysates (see schematic in Fig.1A) and separately measured tissue levels of cAMP using enzyme-linked immunosorbent assay (ELISA). We found that, in fact, PDE4D phosphorylation was doubled in male *Cc2d1a*-deficient hippocampi (2.05±0.13 of WT, p=0.0001 ***, N=5 WT;5 cKO, Fig.1B). cAMP levels were almost halved (WT=51.83±6.88 pmol/mg, N=4; cKO=27.74±1.92 pmol/mg, N=4, p=0.011 *, Fig.1C). Finally, phosphorylation of both Thr-197 in PKA (0.48±0.13 of WT, p=0.019 *, N=4 WT;4 cKO, Fig.1D) and Ser-133 in CREB (0.70±0.09 of WT, p=0.046 *, N=5 WT; 5cKO, Fig.1E) were also decreased. These findings indicate that in the adult hippocampus lacking *Cc2d1a* PDE4D activity is increased leading to baseline reduction in cAMP levels and PKA and CREB phosphorylation which could underlie the cognitive deficits found in behavioral experiments.

**Figure 1.**
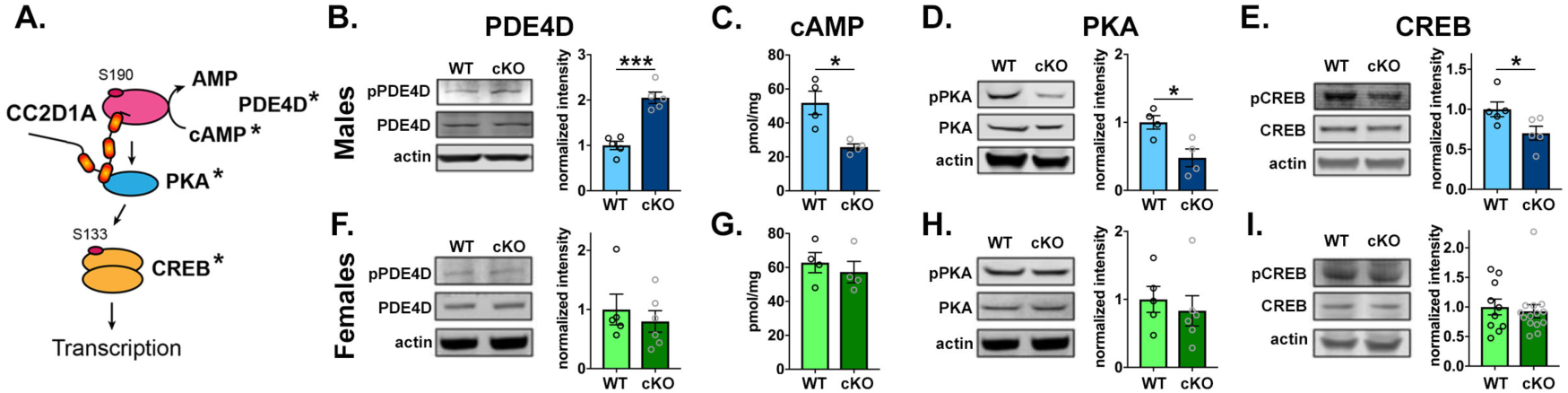
Loss of Cc2d1a causes male-specific disruption of PDE4D and CREB activity in the hippocampus. **A.** Schematic of CC2D1A as a signaling scaffold for PDE4D and PKA upstream of CREB. Expression levels and activity of molecules marked with an asterisk were tested in our studies. **B-E.** Conditional removal of *Cc2d1a* in cKO male mice causes increased phosphorylation and activity of PDE4D (**B.**), leading to reduced levels of cAMP (**C.**), and decreased phosphorylation of both PKA (**D.**) and CREB (**E.**) in hippocampal lysates. **F-I.** No changes in PDE4D (**F.**), cAMP (**G.**), PKA (**H.**) or CREB (**I.**) are found in the female hippocampus. Results expressed as mean ± SEM. Data averages and statistical information are reported in the Results. Two-tailed t-test with equal variance * p<0.05, ** p<0.01, *** p<0.001

Surprisingly, when we tested the same biochemical changes in females, we found that there was no difference in any of them (PDE4D: cKO 0.80±0.45 of WT, p=0.53, N=5 WT; 6 cKO. cAMP: WT=62.85±5.93pmol/mg, cKO=57.28±6.30pmol/mg; p=0.543, N=4 WT; 4 cKO. PKA: cKO 0.83±0.22 of WT, p= 0.59, N=5 WT; 6 cKO. Fig.1F-H). Due to the high variability in CREB activity in female control hippocampi, we increased the number of animals tested to confirm that there was no difference and saw no change (CREB: cKO 0.93±0.11 of WT, p=0.69, N=10 WT;14 cKO. Fig.1I). We tested for similar pathway alterations in cortical lysates where CC2D1A is also removed by CaMKIIa-cre and found that PDE4D phosphorylation and cAMP levels were comparable to WT in cKO males (**Suppl.** Fig.1A-B). As a control, the amount of CC2D1A was measured in the cortex and hippocampus and we found no differences in baseline WT CC2D1A levels between males and females in the two tissues (**Suppl. Fig.1C-D**) and no difference in CC2D1A removal by CaMKIIa-cre mediated recombination (**Suppl. Fig.1E-F**). In summary, we found that loss of *Cc2d1a* causes male-specific activation of PDE4D, leading to reduced CREB activity only in the hippocampus.

**Supplemental Figure 1.**
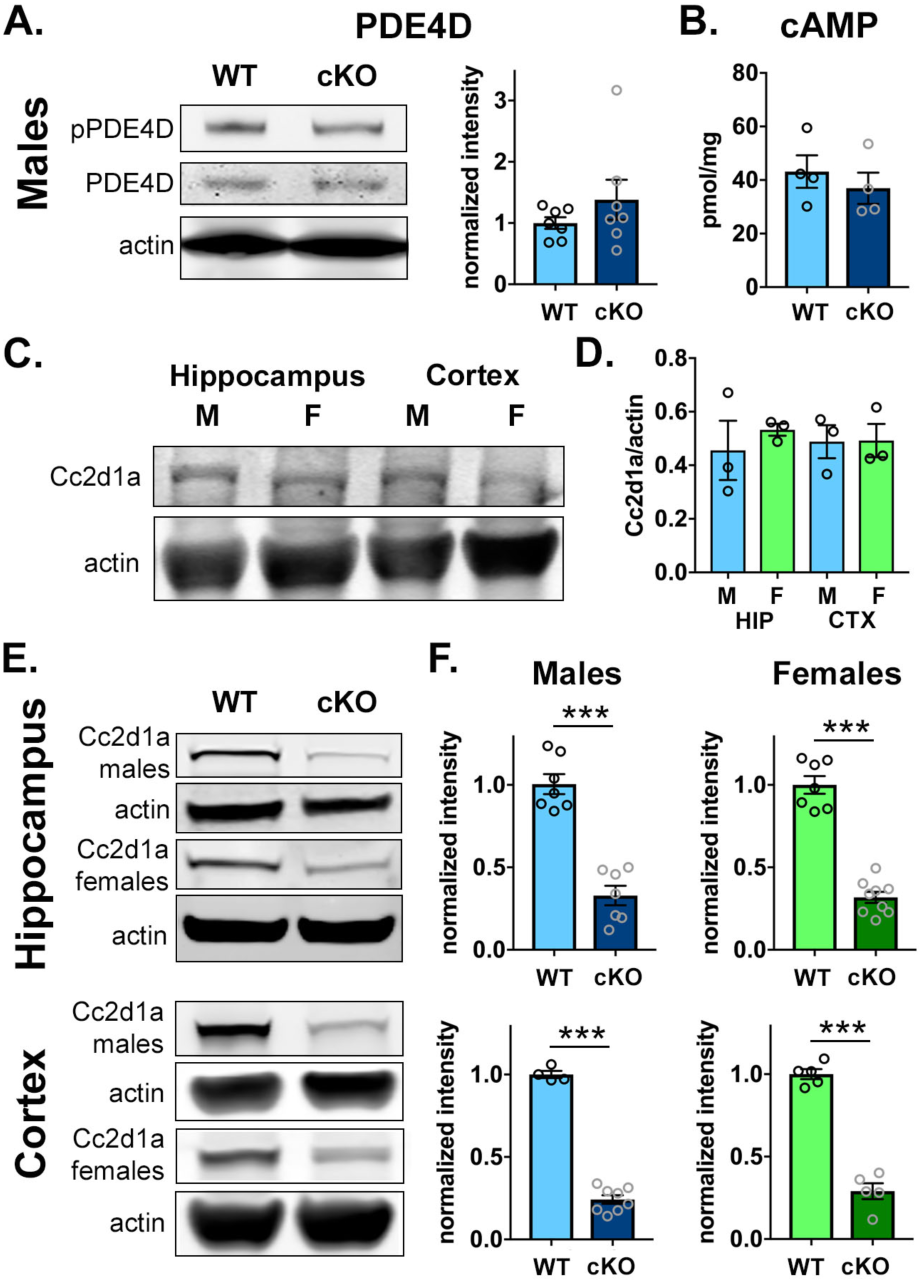
Control studies to test for spatial– and sex-specificity of signaling deficits caused by loss of Cc2d1a A-B. PDE4D phosphorylation measured by Western blot (**A.**) and cAMP levels measured by ELISA (**B.**) are not changed in the cortex of *Cc2d1a* cKO males indicating that the signaling changes shown in Fig.1 are specific to the hippocampus. **C-D.** CC2D1A expression levels are the same in male and female WT brains in the hippocampus and cortex (quantified in D.). **E-F.** Reduction in CC2D1A expression is comparable in males and females in the cortex and hippocampus showing that sex-specific signaling differences are not due to residual CC2D1A activity in the female hippocampus. Note that CC2D1A expression is not completely ablated because the CaMKIIa promoter is primarily active in excitatory neurons causing conditional CC2D1A removal only in that population and leaving CC2D1A in interneurons intact.

### Cc2d1a cKO females only show cognitive deficits in object recognition

*Cc2d1a* cKO males show an array of behavioral deficits: a failure to discriminate between known and a novel object in the Novel Object Recognition Test (NORT), a defect in spatial memory shown by a delay in memorizing the location of hidden platform in the Morris Water Maze (MWM), reduced sociability in the three-cambered test, and increased activity and reduced time in center in the Open Field test (OF) (20). We tested cKO females through the same battery of tests used for the males, after general somatosensory and motor function testing which was normal (**Suppl.** Table 1).

**Table 1.**
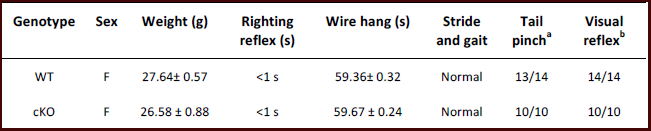
Analysis of basic motor and somatosensory function

In the NORT animals are trained on two identical object and after a short stay in their home cage they are challenged with one of the previous objects and a novel object. Mice are neophilic and will spend more time exploring the novel object (WT training ratio=0.76±0.09, WT novel ratio=2.08±0.44, p=0.0063 **, N=18, Fig.2A). Similarly to cKO males, cKO females did not recognize the novel object (cKO training ratio=1.41±0.18, cKO novel ratio= 1.53±0.23, p=0.71, N=13, Fig.2A) and showed reduced interest in the objects, as shown by significantly reduced overall exploration time, and reduced exploration in the novel object trial (Training trial: WT=54.01±8.01s, cKO=38.64±7.91s, p=0.195; Novel trial: WT=63.73±12.16s, cKO=30.45±6.50, p=0.023 *; Total: WT=117.7±12.99s, cKO=69.09±12.94s, p=0.015 *, N=18 WT, 13 cKO. Fig.2B).

**Figure 2.**
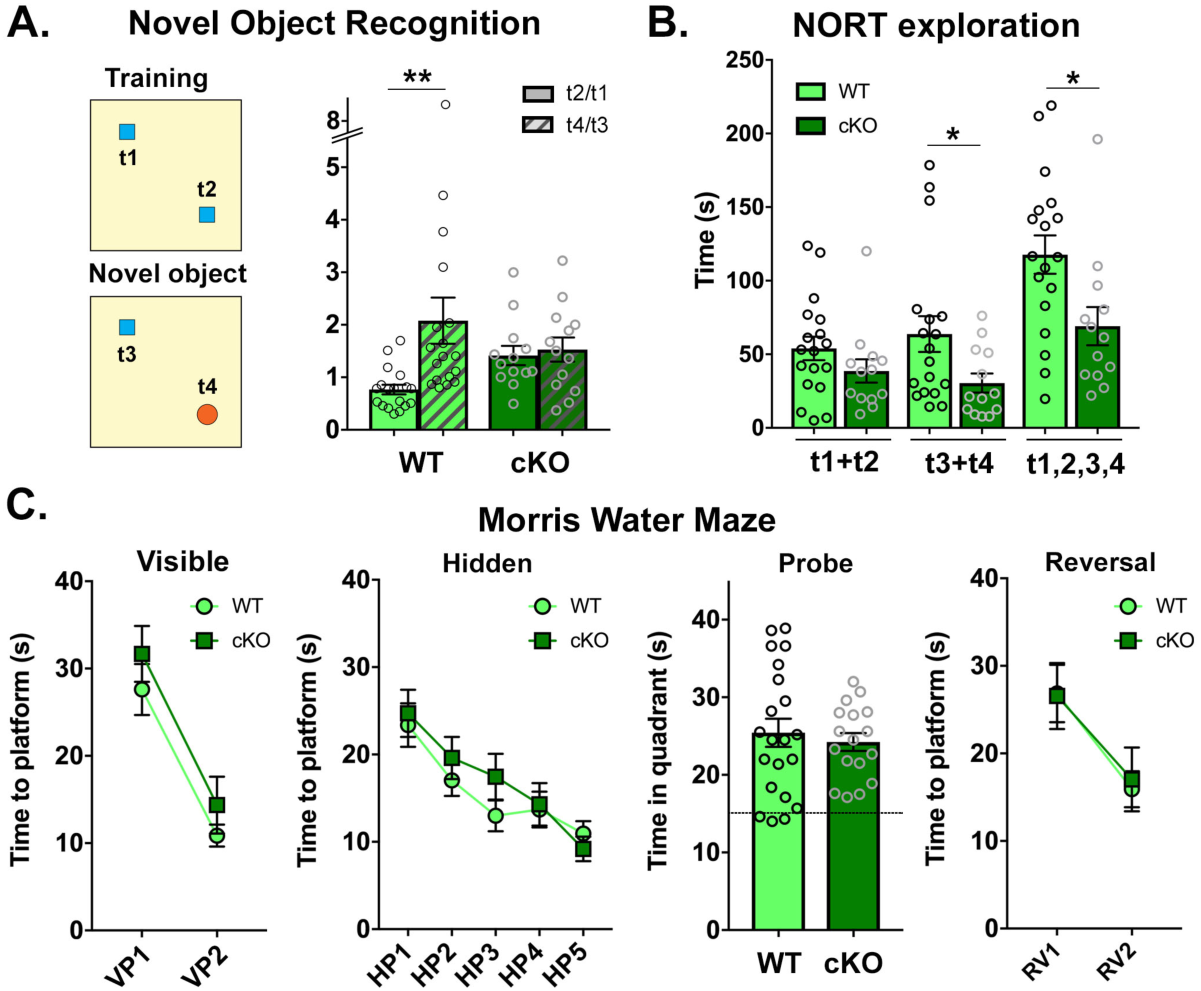
*Cc2d1a* cKO females show a deficit in object memory, but not in spatial memory. **A.** Left. Schematic design of short-term novel object recognition test (NORT), with a novel object replacing a familiar object after a 15 min interval. Right. In contrast to WT, cKO female mice did not show a preference for the novel object relative to a familiar object. Results expressed as mean ± SEM, ** p<0.01 (two-tailed t-test with equal variance). **B.** Summary of exploration times is divided by initial exploration of training objects 1 and 2 (t1+t2), test objects 3 and 4 (t3+t4) and total exploration across the two phases of the NORT. cKO females show significantly reduced exploration during the test (t3+t4), and when total exploration is calculated as the sum of t1,2,3,4. Results expressed as mean ± SEM. Two-tailed t-test with equal variance, * p<0.05. **C.** Hippocampus-dependent spatial memory was assessed in cKO females via the Morris Water Maze test. Spatial learning was measured as latency to reach the platform in three different stages, visible platform (VP) and hidden platform (HP). Then, spatial memory retention was measured after the HP trial in the Probe trial by showing that the time spent swimming in the quadrant where the platform was previously located was increased above chance (dotted line). Finally, acquisition of a different platform location was tested in the reversal (RV) trial. Differently from cKO males (20) which show a delay in the HP trial, cKO females showed no deficit.

The same cohort was also tested in the Morris Water Maze (MWM) spatial memory test, where mice must use cues on the walls of a tub filled with water to memorize the location of a platform hidden under the surface. After an initial trial with a visible platform (VP) to learn that a platform is available to escape the water, each mouse is tested in multiple trials with a hidden platform (HP). The platform is then removed to test for permanence of the spatial memory (probe) and moved to a different location (reversal trial, RV) to asses cognitive flexibility (27,30). Cc2d1a cKO males had showed a defect in learning the HP location in days 2 and 3 of the 5 day hidden test, but had normal performance in the probe and RV test (20). cKO females (N=21WT, 17cKO) showed equal performance to controls in all phases of the test and demonstrated normal spatial memory (Fig.2C). While sex has been shown not to affect performance in the MWM test in C57BL/6 mice (31), studies in rats showed that males and females can use different strategies to acquire the position of the hidden platform shifting from place-based strategies using visual cues, to motor-based search strategies such as scanning the length of tub (9). To confirm that females were not just using a different strategy to identify the location of the platform we compared learning strategies during the first three days of the HP test with our published male cKO cohort, using criteria outlined by Garthe et al (32). WT males and females showed no differences in using motor- (thigmotaxis, random search or scanning), and place-based strategies (circling/chaining, directed search, or straight line) (**Suppl.** Fig.2A). cKO males showed a trend towards being slower in transitioning to place-based strategies in day 2 and 3 (**Suppl.** Fig2D-G). The primary factor driving the difference in performance in cKO males was the latency from drop location to the platform.

cKO females also did not show any difference from control females in the open field or the three-chambered test (Fig.3). While cKO males showed significant hyperactivity and avoided the center of the open field arena (20), females of both genotypes traveled equal distances (WT=35.34±3.32m, N=14; cKO=33.49±4.76m, N=10; Fig.3A) and spent similar amounts of time in the center of the apparatus (WT=62.66±11.05s, N=14; cKO=52.50±10.53s, N=10; Fig.3A). Similarly, while cKO males showed reduced sociability (20), female cKOs were equal to control females in preferring a novel mouse (WT=352.30±20.79s, N=8; cKO=345.34±26.54s, N=8; Fig.3B) to an empty enclosure (WT=161.13±15.89s, N=8; cKO=173.09±25.70s, N=8; Fig.3B). The only other phenotype which was found in both cKO females and males was ulcerative dermatitis, which appeared in females at a rate of 33.3% (N=16/48) starting after 3-4 months of age, while WT females only showed dermatitis in 2.7% of animals (N=1/37).

**Figure 3.**
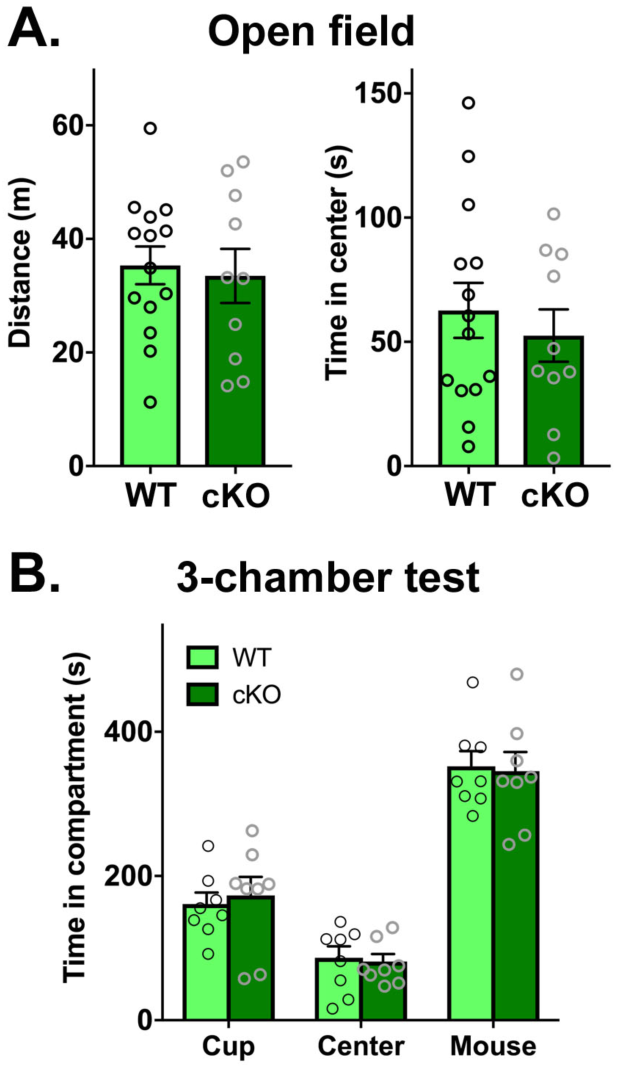
Cc2d1a cKO females show no deficits in the open field or three-chambered test. **A.** Exploratory and general locomotor activity in a novel environment were assessed on the open field test. Differently from Cc2d1a cKO males which had shown increased locomotor activity and decreased time in center in the in the Open Field test, cKO females were comparable to WT lit-termates. **B.** Social interaction behavior was assessed by the three-chamber test, presented as time spent in each chamber: empty center, with novel mouse contained in a metal enclosure (mouse) or with an empty enclosure (cup). *Cc2d1a* cKO females showed WT-like sociability. Re– sults expressed as mean ± SEM. Data averages and statistical information are reported in the Results.

Overall, these findings show that *Cc2d1a* cKO females are less severely affected than males, as they only display a deficit in object recognition in the NORT. They have normal spatial memory, sociability and activity levels, indicating that loss of *Cc2d1a* also causes sex-specific behavioral deficits in addition to sex-specific signaling changes.

**Supplemental Figure 2.**
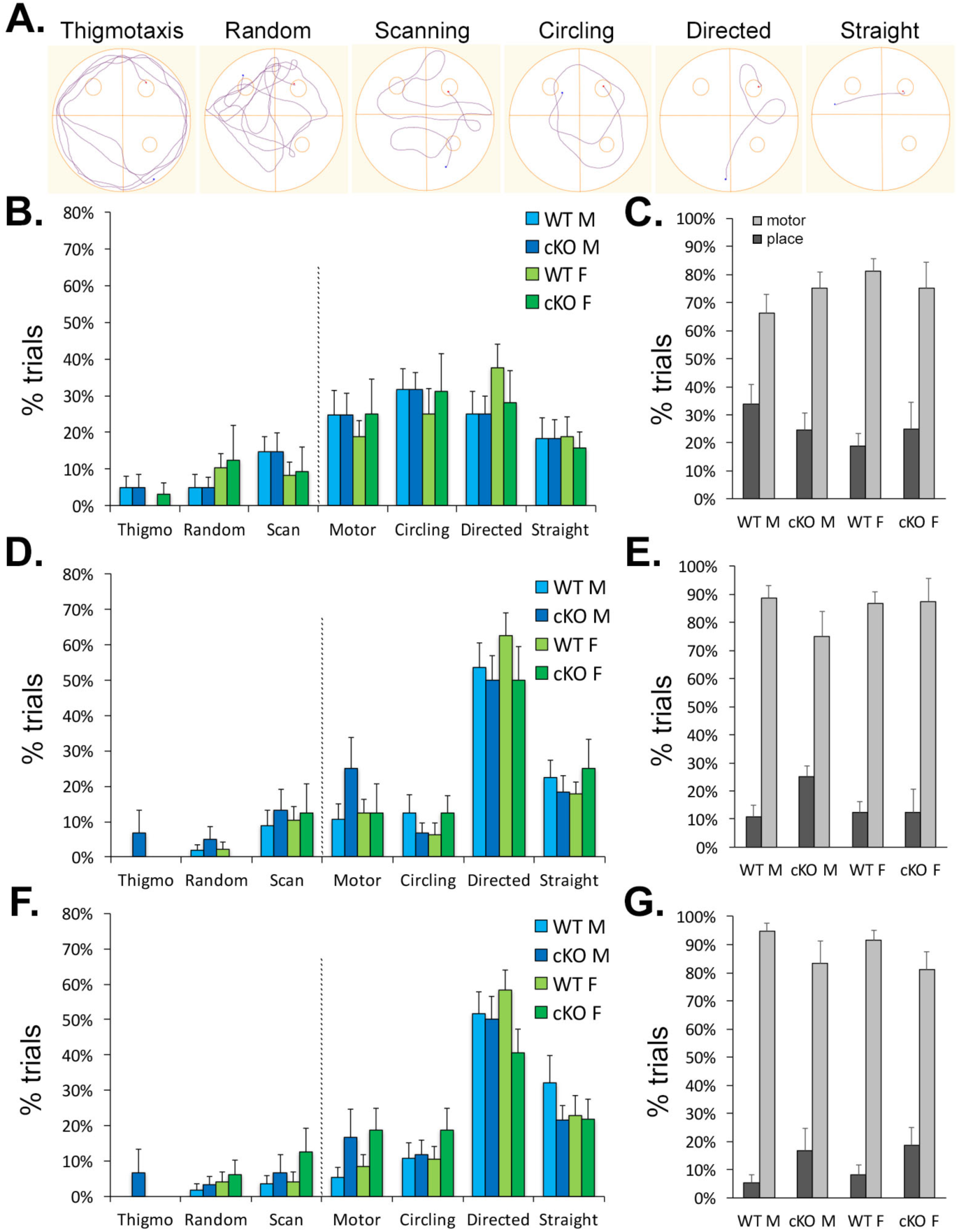
Analysis of learning strategies in the Morris Water Maze test. A. Six different learning strategies were assayed using criteria from Garthe et al (ref.32). Thigmotaxis, random path and scanning were grouped as motor strategies where the animal does not appear to use visual cues to locate the platform. Circling around the tub at the distance of the platform or swimming toward the platform with (directed) or without errors (straight) indicate that the animal is using a place-based strategy. B-G. Average use of different strategies is broken down for males (M) and females (F) showing each strategy separately and motor- vs. place-based strategies for day 1 (**B-C.**), day 2 (**D-E.**) and day 3 (**F-G.**).

### Adult restoration of PDE4D activity and CREB signaling rescues spatial memory deficits in Cc2d1a cKO males

CREB activity is critical for creating and storing spatial memories (13,14,33) and its activation levels must be carefully controlled (34). Upstream of CREB, PDE4D is also a key regulator of cognitive function (35). We asked whether inhibiting PDE4D would reduce cAMP degradation to restore control levels of cAMP and CREB activity this rescuing spatial memory deficits in male cKOs (Fig.4A). PDE4 inhibitors have been studied in dementia and Alzheimer’s Disease to slow cognitive decline by enhancing CREB function (36,37). However, the most commonly used drug, rolipram, targets multiple PDE4s and leads to severe side effects, including nausea and emesis which cause poor tolerability (36). To inhibit PDE4D in the cKO hippocampus we used a specific inhibitor GEBR-7b, which is selective for PDE4D and better tolerated (38). Concentrations of 30 µg/kg of GEBR-7b had proven effective in improving spatial memory in both WT mice and a mouse model of Alzheimer’s Disease (38,39), and were used for a 14-day course of treatment via intraperitoneal injections in a cohort of adult control and cKO male and female mice. Treatment was effective at restoring levels of cAMP as tested by ELISA (WT saline=47.17±2.97 pmol/mg, N=4, cKO saline=29.33±2.69 pmol/mg, N=4, p=0.014 to WT; WT GEBR=46.37±3.02 pmol/mg, N=4, cKO GEBR=47.97±4.91 pmol/mg, N=3. Fig.4B) and levels of CREB phosphorylation as tested by Western blot (cKO saline=0.47±0.08 of WT saline, p=0.048 *, N=7 WT, 6 cKO; WT GEBR=1.19±0.26, N=6; cKO GEBR=0.82±0.07, N=7. Fig.4C).

**Figure 4.**
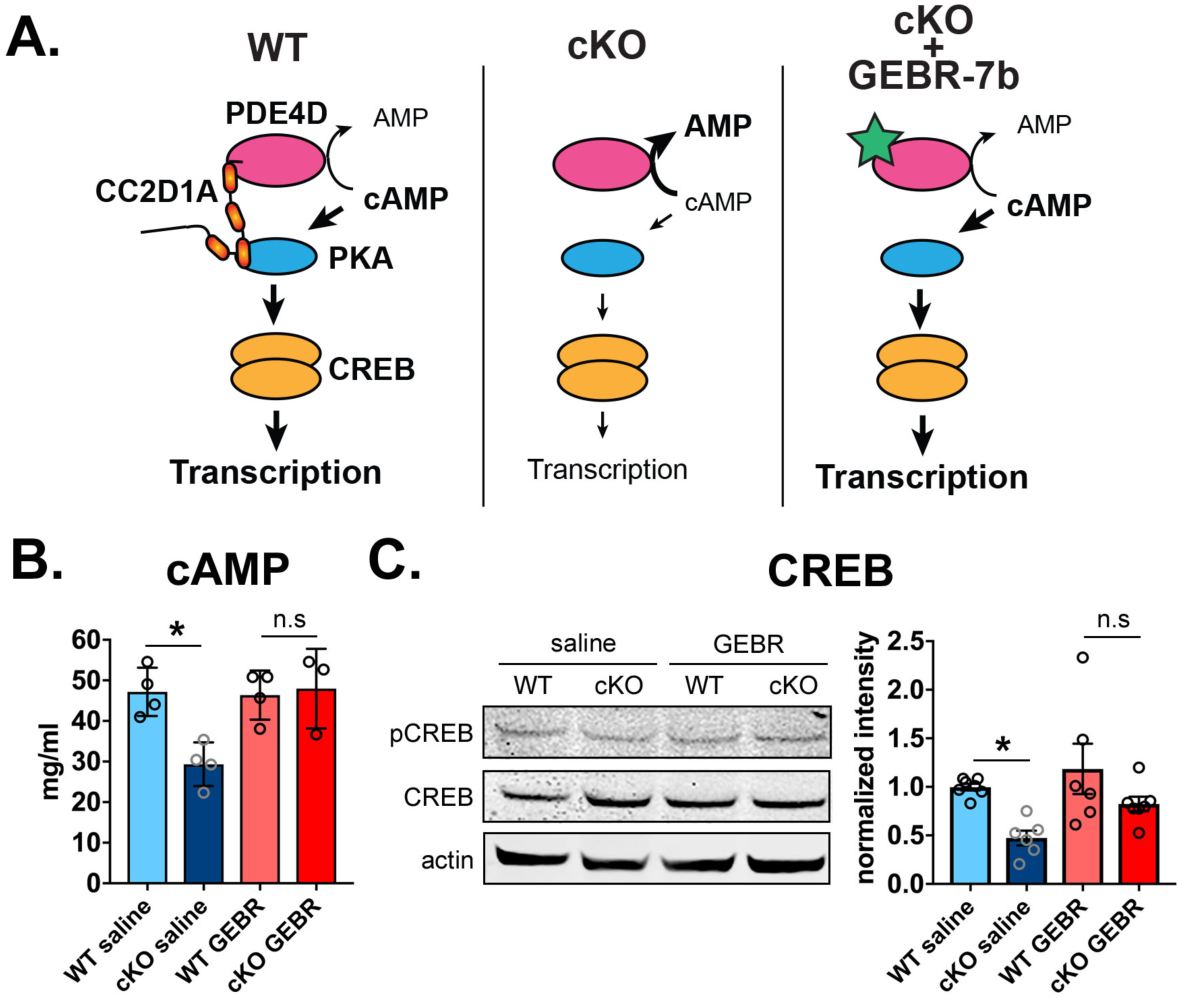
GEBR-7b treatment restores cAMP levels and CREB phosphorylation in *Cc2d1a* cKO males. A.s. Schematic of expected action of GEBR-7b (green star). When CC2D1A is absent hyperactivity of PDE4D in cKO males increases cAMP degradation to AMP (center panel). GEBR-7b is expected to restore cAMP levels and CREB-mediated transcription (right panel). **B.** 14-day daily treatment with 30µg/kg of GEBR-7b (GEBR) restored cAMP to WT levels in the hippocampus of cKO males when compared to animals treated with saline. **C.** CREB phosphorylation is also increased in GEBR-7b treated cKO males. One-way ANOVA with Tukey’s multiple comparison test, *p<0.05.

After treatment, the mice were tested in an abbreviated MWM protocol consisting of the visible, hidden, and probe tests, since neither males nor female cKO has shown a difference in reversal test. In this cohort, cKO males again showed a delay in acquiring the position of the hidden platform when compared to controls in day 2 and the deficit was completely rescued by GEBR-7b treatment (HP2: WT saline=9.86±1.51s, N=12; cKO saline=25.38±4.88s, N=9; WT GEBR=11.80±1.94s, N=12; cKO GEBR=8.29±1.41, N=10; p values to cKO saline: WT saline p=0.0044 **, WT GEBR p=0.015 *, cKO GEBR p=0.0037 ** Fig.5A). Following spatial learning treated and untreated males performed equally in the probe trial (Fig.5B), and showed no differences in swim speeds showing that treatment did not change motor function (Fig.5C). Females performed equally with and without GEBR-7b treatment (Fig.5D). No significant differences were found in probe test performance (Fig.5E) or in swim speed (Fig.5F). Overall our results show that CC2D1A established male-specific CREB signaling through PDE4D that controls spatial memory acquisition.

**Figure 5.**
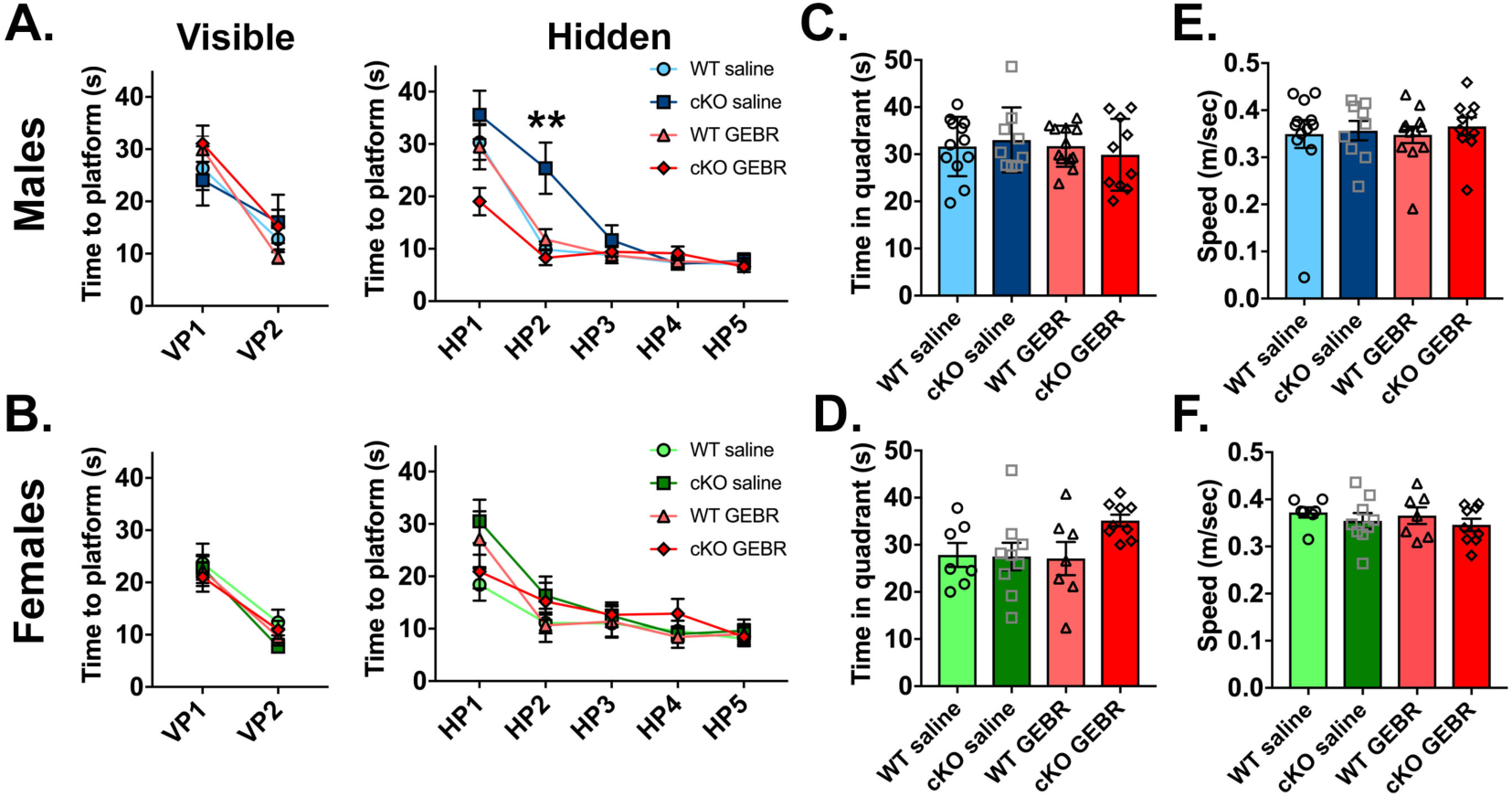
GEBR-7b treatment rescues spatial memory deficit in Cc2d1a cKO males and has no effect on females. **A.** The learning delay in the hidden platform (HP) trial of the MWM found in male Cc2d1a cKO animals is corrected by 30µg/kg GEBR-7b treatment for 14 days before testing. Multiple comparison t-tests at HP2, ** p<0.01 between WT saline and cKO saline. **B.** GEBR-7b treatment has no effect on females. **C-D.** No significant change is found in the Probe test where all genotypes and treatments perform above chance (dotted line). cKO females treated with GEBR-7b show a trend for better performance in this test. **E-F.** GEBR-7b treatment did not affect swim speed.

## DISCUSSION

Here, we demonstrated that the ID/ASD gene *CC2D1A* controls male-specific CREB signaling in the hippocampus leading to male-specific spatial memory deficits in mice. We found that loss of CC2D1A in males increased hippocampal PDE4D phosphorylation leading to a reduction of cAMP levels and of CREB phosphorylation which can be rescued by PDE4D inhibition in cKO males. cKO female mice, however, do not show any of the signaling deficits observed in males and are not impaired in hidden platform acquisition in the MWM. Our findings identify a novel molecular mechanism for sex-specific signaling regulation controlled by a single gene mutated in ID and ASD.

Despite equal loss of CC2D1A expression in male and female cKO mice, females only present with a subset of behavioral deficits found in males, novel object recognition deficits and obsessive grooming leading to ulcerative dermatitis, but do not display the spatial deficits, hyperactivity, and anxiety found in cKO males. Multiple hypotheses have been put forth to explain the male bias in neurodevelopmental disorders (40,41). Increased levels of steroid hormones, including cortisol and testosterone, and their precursors have been found elevated in the amniotic fluid of pregnancies that resulted in individuals with ASD (42), suggesting a role for sex hormones and other steroids in the etiology of the disorder in males. On the other hand, females require more severe rare genetic mutations and more familial risk factors than males to become affected (43,44), supporting the hypothesis that females are protected. Our findings in *Cc2d1a* cKO mice, together with those from Grissom et al in a 16p11.2 hemideletion model (15), reveal that sex-specific disruptions in intracellular signaling may be involved in establishing male-specific behavioral phenotypes. While these findings do not yet define whether males are susceptible or females are protected, they show that in inbred genetic models recapitulating human mutations equally in males and females, males develop sex-specific signaling deficits. Defining how these signaling differences are established will not only provide insight into the mechanisms of sex bias in the pathogenesis of neurodevelopmental disorders, but will also likely identify sex-specific targets for therapy development.

Our studies show that PDE4D inhibition can rescue cognitive deficits in a model of neurodevelopmental disorders. PDE4 proteins have been under investigation as a target for the improvement of cognitive function in neurodegenerative disorders for some time, since CREB activity is disrupted in multiple models of Alzheimer’s disease (AD) (36,37,45). However, the most commonly used pan-PDE4 inhibitor rolipram is poorly tolerated in humans because of severe emesis triggered by PDE4D activity (36). Since PDE4D is highly expressed in the cortex and hippocampus (46) and has a critical role in cognitive function (47), multiple selective PDE4D inhibitors acting at sub-emetic concentrations are being developed (48). GEBR-7b is highly selective for PDE4D (49) and improved cognitive performance in both WT mice and mouse models of AD at non-emetic doses (38,39). It will be important to determine whether PDE4D function is disrupted in other models of neurodevelopmental disease, and whether PDE4D inhibition could be beneficial in ID. It should also be noted that AD and dementia show a strong sex-bias which is opposite than the bias observed in ID and ASD, with females comprising two thirds of individuals affected by AD (50). Sex differences in the regulation of PDE4D and CREB signaling at different ends of the human life-span may need to be considered in developing therapies.

To date, *CC2D1A* mutations have been identified in 34 individuals (21 males and 13 females) with fully penetrant developmental delay and ID in males and females, and partially penetrant reports of ASD, ADHD, aggressive behavior, and seizures, primarily in males (16-19). Since most of the individuals with CC2D1A mutations reside in the Middle East and are not currently accessible to follow-up, we do not know whether there are significant differences in the behavioral presentation and severity between males and females. Intriguing sex-specific differences in *CC2D1A* expression were found in Major Depressive Disorder (MDD). Conditional *Cc2d1a* removal in serotoninergic neurons in the dorsal raphe demonstrated that CC2D1A also has a role in regulating serotonin receptor function and depression-like behaviors (51). *CC2D1A* levels were found reduced in the prefrontal cortex (52) and increased in the cingulate cortex (53) in males, but not females affected by MDD, suggesting that male-specific impairment in CC2D1A function in humans could be present. *CC2D1A* expression was elevated in the blood of individuals with ASD (54), but the number of females in the sample was too small to explore sex-specificity of this change and more extensive follow-up will be necessary. PDE4D expression is also regulated by both serotonin and norepinephrine reuptake inhibitors (46,55), suggesting a shared role for CC2D1A and PDE4D in cognition and depression.

CC2D1A has also been studied in regulation of subcellular and endosomal regulation of signaling proteins (22,56,57) and future investigations should focus on whether this function is regulated by sex hormones and/or involved in sex hormone function. Defining how these mechanisms control cognitive and affective function throughout life may provide insight into sex bias in disease susceptibility.

## ACKNOWLEDGEMENTS

The authors are grateful to the Animal Research Facility at GWU SMHS, Judy Liu, Maria Chahrour, Ottavio Arancio, and Emanuela Santini for helpful discussion on experimental design. This work was supported by start-up funds from the George Washington University (M.C.M.), NIH grant R00HD067379 (M.C.M.) and Pilot Grants (M.C.M) from the Intellectual and Developmental Disabilities Research Center (IDDRC) at Children’s Research Institute (P30HD040677) and from the Clinical and Translational Studies Institute at Children’s National (UL1TR001876). The *Cc2d1a* KO mouse strain used for this research project was generated by the trans-NIH Knock-Out Mouse Project (KOMP) and obtained from the KOMP Repository (www.komp.org). NIH grants to Velocigene at Regeneron Inc (U01HG004085) and the CSD Consortium (U01HG004080) funded the generation of gene-targeted ES cells for 8500 genes in the KOMP Program and archived and distributed by the KOMP Repository at UC Davis and CHORI (U42RR024244). For more information or to obtain KOMP products go to www.komp.org or.

